# MAGI-MS: Multiple seed-centric module discovery

**DOI:** 10.1101/2021.09.14.460296

**Authors:** Julie C. Chow, Ryan Zhou, Fereydoun Hormozdiari

## Abstract

Complex disorders manifest by the interaction of multiple genetic and environmental factors. Through the construction of genetic modules that consist of highly co-expressed genes, it is possible to identify genes that participate in common biological pathways relevant to specific phenotypes. We have previously developed tools MAGI and MAGI-S for genetic module discovery by incorporating co-expression and protein-interaction networks. Here we introduce an extension to MAGI-S, denoted as Merging Affected Genes into Integrated Networks - Multiple Seeds (MAGI-MS), that permits the user to further specify a disease pathway of interest by selecting multiple seed genes likely to function in the same molecular mechanism. By providing MAGI-MS with pairs of seed genes involved in processes underlying certain classes of neurodevelopmental disorders, such as epilepsy, we demonstrate that MAGI-MS can reveal modules enriched in genes relevant to chemical synaptic transmission, glutamatergic synapse, and other functions associated with the provided seed genes.

**Availability and implementation:** MAGI-MS is free and is available at: https://github.com/jchow32/MAGI-MS

## Introduction

The extensive genetic and phenotypic heterogeneity characteristic of complex disorders indicates that the interaction of multiple genes underlie etiology (Parenti *et al*., 2020). The development of protein-protein interaction and co-expression networks has aided in identification of networks of genes hypothesized to belong to the same functional module and contribute to specific pathways (Parikshak *et al*., 2015; Chen *et al*., 2020).

Previously, we described a method called MAGI-S used to dissect complex phenotypes, such as epilepsy, by producing modules seeded from a single gene associated with the phenotype of interest (Chow *et al*., 2019). We demonstrated that independently providing MAGI-S single seed neurodevelopmental disorder (NDD) genes with varying degrees of association with epilepsy revealed modules enriched in 1) non-synonymous coding *de novo* variation in affected NDD cases relative to controls, 2) genes associated with epilepsy, and 3) *de novo* mutation specifically retrieved from epilepsy cohorts, suggesting that MAGI-S can uncover networks of genes relevant to a complex disorder.

We introduce an extension to the existing method MAGI-S (Chow *et al*., 2019), referred to as MAGI-Multiple Seeds (MAGI-MS). MAGI-MS permits the user to select up to three seed genes from which to construct modules, using either the average or minimum co-expression of other genes relative to the selected seeds during gene score assignment. In addition, we have simplified the process of running the compiled MAGI-MS program by providing example commands, sample input files, and suggested parameter combinations for ease of use.

## Materials and methods

MAGI-MS uses a protein-protein interaction (PPI) network, co-expression network, deleterious mutations within a control population, and seed gene(s) to create genetic modules that satisfy constraints related to PPI connectivity and degree of co-expression amongst module genes (**Supplementary Data**). In the following experiments, we use PPIs retrieved from the HPRD and the STRING databases (Keshava Prasad *et al*., 2009; Szklarczyk *et al*., 2011), RNA-seq data from the BrainSpan: Atlas of the Developing Human Brain as the co-expression network (Miller *et al*., 2014), and control variants from the NHLBI Exome Sequencing Project (ESP) (http://evs.gs.washington.edu/EVS/). Briefly, MAGI-MS assigns a score (**Equation 1**) to every gene within the PPI network (**Figure 1**). High-scoring seed pathways are created by the use of a modified color coding algorithm to find simple paths that maximize the summation of scores associated with genes (Hormozdiari *et al*., 2015). Seed pathways are then merged into clusters by a random walk, and clusters are improved incrementally by local search to yield top scoring modules.

**Figure 1.**
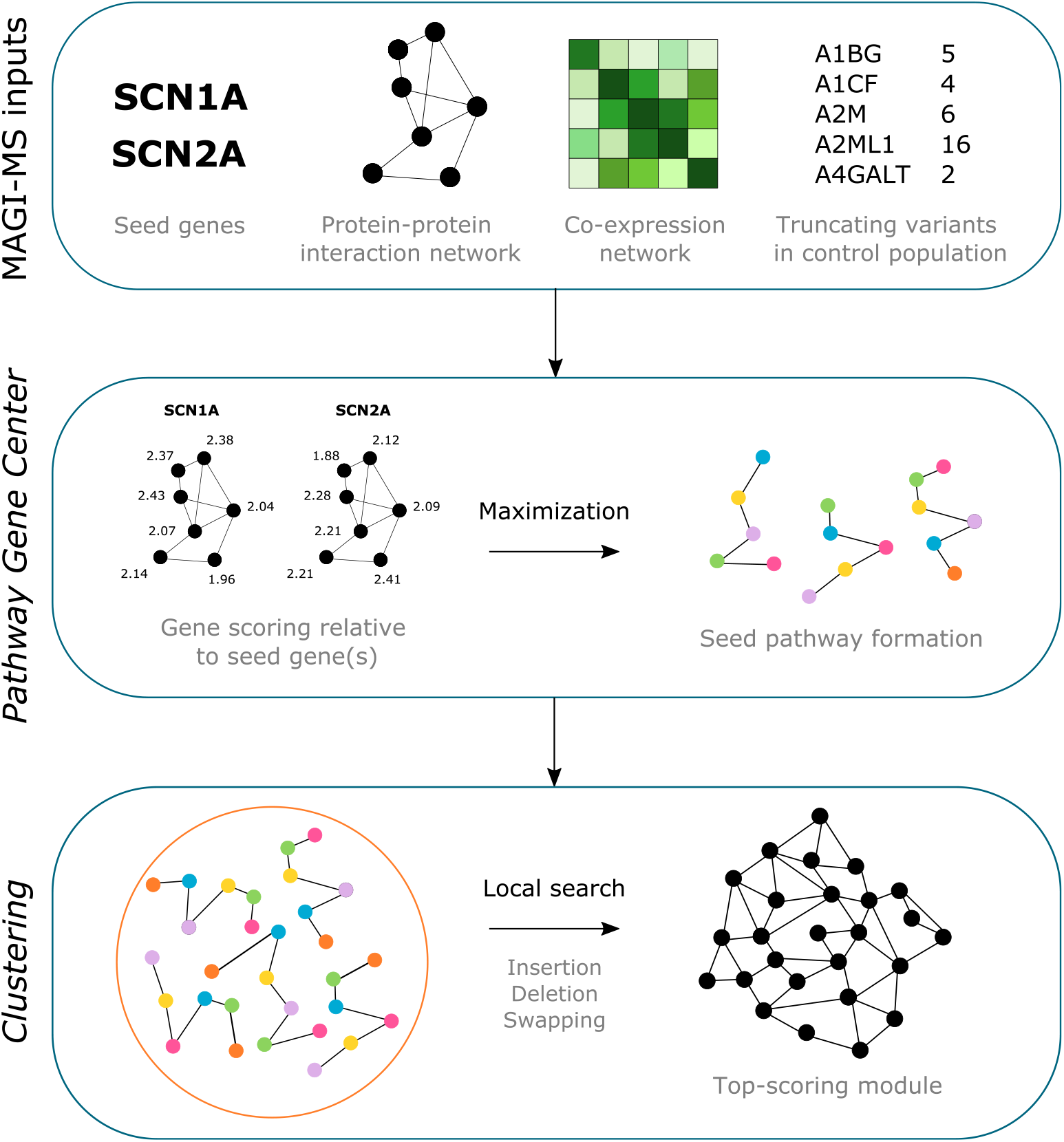
General methods overview of MAGI-MS. User-selected seed gene(s), a protein-protein interaction (PPI) network, a co-expression network, and loss-of-function mutations observed in a control population are provided as input to construct modules specific to biological pathways associated with the provided seed genes. During *Pathway Gene Center*, scores are assigned to genes to describe their degree of co-expression with seed gene(s), and seed pathways consisting of high scoring genes are formed. During *Clustering*, seed pathways are merged and refined to produce candidate modules.

To assess the ability of MAGI-MS to dissect a complex phenotype, we provided MAGI-MS with 6 pairs of seed genes, where each pair consists of genes observed to participate in a similar biological function (Szklarczyk *et al*., 2021) (CHD8-CREBBP, CHD8-CTNNB1, GABRA3-GABRB1, GRIN2A-GRIN2B, SHANK2-SHANK3, SCN1A-SCN2A): We additionally provided MAGI-MS with a seed gene pair that are not hypothesized to participate in the same pathways (SCN1A-CTNNB1). To confirm the presence of relevant functional enrichment and cell-type specific expression, modules were provided to the tools Enrichr and Cell-type Specific Expression Analysis (CSEA) (Kuleshov *et al*., 2016; Xu *et al*., 2014).

## Results

Given pairs of seed genes involved in the same biological pathway, MAGI-MS produces modules that have significant overlap with modules seeded from either seed gene alone (**Supplementary Table 1**). On average, 49.5% and 61.4% of the genes in paired modules exist, using either minimum or average co-expression values during gene score assignment, respectively, in either of the singly-seeded modules.

Modules with paired seeds related to the epilepsy phenotype (GABRA3-GABRB1, GRIN2A-GRIN2B, and SCN1A-SCN2A) were enriched in terms such as long-term potentiation, chemical synaptic transmission, among others, and showed selective expression in deep cortical neurons (**Supplementary Table 1**). For seed gene pairs related to more general NDD and autism phenotypes (CHD8-CREBBP and CHD8-CTNNB1), we observe an enrichment in chromatin organization and regulation of transcription. For the module constructed with seed genes that do not participate in the same biological function (SCN1A-CTNNB1), a module was not formed due to low scoring seed pathways, indicating that the choice of multiple seed genes from pathways with similar biological function is critical to form a module that is useful for the dissection of a specific phenotype.

## Conclusion

We present an extension to the existing method MAGI-S, denoted as MAGI-MS, which permits the discovery of genetic modules specific to certain biological functions by selection of multiple seed genes involved in a pathway of interest. MAGI-MS is freely available with updated user guides for parameter and input choices.

## Supplementary data

SupplementaryTable1 - .xlsx file

SupplementaryData - .PDF file

## References

Chen, S. et al. (2020) De novo missense variants disrupting protein–protein interactions affect risk for autism through gene co-expression and protein networks in neuronal cell types. Mol. Autism, 11, 76.

Chow, J. et al. (2019) Dissecting the genetic basis of comorbid epilepsy phenotypes in neurodevelopmental disorders. Genome Med., 11, 65.

Hormozdiari, F. et al. (2015) The discovery of integrated gene networks for autism and related disorders. Genome Res., 25, 142–154.

Keshava Prasad, T.S. et al. (2009) Human Protein Reference Database—2009 update. Nucleic Acids Res., 37, D767–D772.

Kuleshov, M.V. et al. (2016) Enrichr: a comprehensive gene set enrichment analysis web server 2016 update. Nucleic Acids Res., 44, W90–W97.

Miller, J.A. et al. (2014) Transcriptional landscape of the prenatal human brain. Nature, 508, 199–206.

Parenti, I. et al. (2020) Neurodevelopmental Disorders: From Genetics to Functional Pathways. Trends Neurosci., 43, 608–621.

Parikshak, N.N. et al. (2015) Systems biology and gene networks in neurodevelopmental and neurodegenerative disorders. Nat. Rev. Genet., 16, 441–458.

Szklarczyk, D. et al. (2011) The STRING database in 2011: functional interaction networks of proteins, globally integrated and scored. Nucleic Acids Res., 39, D561–D568.

Szklarczyk, D. et al. (2021) The STRING database in 2021: customizable protein-protein networks, and functional characterization of user-uploaded gene/measurement sets. Nucleic Acids Res., 49, D605–D612.

Xu, X. et al. (2014) Cell Type-Specific Expression Analysis to Identify Putative Cellular Mechanisms for Neurogenetic Disorders. J. Neurosci., 34, 1420–1431.

