## Supplementary Data for "MAGI-MS: Multiple seed-centric module discovery"

**Supplementary methods**

MAGI-MS takes as input a protein-protein interaction (PPI) network, a co-expression network, loss-of-function mutation from control populations, and up to three user-selected seed gene(s). The PPI network was retrieved from HPRD and STRING (Keshava Prasad *et al.*, 2009; Szklarczyk *et al.*, 2011). Normalized RPKM values were retrieved from the BrainSpan: Atlas of the Developing Human Brain (Miller *et al.*, 2014) (V6). Truncated variants from the NHLBI Exome Sequencing Project (ESP) (http://evs.gs.washington.edu/EVS/) were used.

Genes within modules constructed by MAGI-MS must satisfy constraints related to 1) degree of connectivity in the PPI network, 2) high pairwise co-expression among module genes, and 3) restriction of the number of deleterious loss-of-function mutations in module genes from a control population as described in the Supplementary Material of MAGI (Hormozdiari *et al.*, 2015).

*Pathway Gene Center*

To construct modules, first, a score is assigned to every gene within the PPI network denoting its degree of co-expression with the seed gene(s). This score ($G_s$) (**Equation 1**), as in MAGI-S (Chow *et al.*, 2019), is calculated as follows, for every gene *G*:

$$G_s = ((H_1)(H_2)) / N^2) \qquad \textbf{Equation 1}$$

where $H_1$ is equal to the number of pairwise comparisons of co-expression values for which the co-expression value of the [seed gene and the gene to be scored] is greater than the value of the [seed gene and another gene in the PPI network (iteratively through all other genes in the network)], and $H_2$ is equal to the number of pairwise comparisons of co-expression values for which the co-expression value of the [seed gene and the gene to be scored] is greater than the value of the [gene to be scored and another gene in the PPI network (iteratively, as before)]. *N* is the total number of genes within the PPI network.

Gene scores are calculated for every gene relative to each seed gene. For example, if two seed genes are provided, then any particular gene will have two scores, where each score is associated with a different seed gene. For each seed gene *k*, individual gene scores are z-scored (**Equation 2**), such that the scores for any particular seed gene possess a mean score of 0 with standard deviation of 1.

$$z_k = (G_{s,k} - \mu_k) / \sigma_k \qquad \textbf{Equation 2}$$

Following z-scoring, for every gene in the PPI network, a final score is assigned by either taking the average (-avg) or minimum (-min) score among candidate scores from each seed. A larger score indicates a greater degree of co-expression with the seed genes. By calculating an average score, final gene scores will reflect the average degree of co-expression that the gene possesses with seed genes. By using a minimum score, final gene scores will reflect the largest degree of co-expression observed with any of the seed genes provided.

After final scores have been assigned to every gene in the PPI network, seed pathways are formed via the color coding algorithm modified to limit the number of deleterious mutations observed in a control population (Hormozdiari *et al.*, 2015; Alon *et al.*, 1995). Seed pathways consist of *h* genes, in which MAGI-MS seeks to maximize the summation of scores within the seed pathway by randomly coloring genes with *h* different colors and finding the colorful path via dynamic programming. *Pathway Gene Center* produces a total of 16,000 seed pathways using 1,000 iterations of combinations of number of loss-of-function mutations allowed in the control population (0, 1, 2, 3) and number of genes within seed pathways ($h$ = (5, 6, 7, 8)), written to 16 files (*BestPaths* files).

*Clustering*

During the clustering process, seed pathways generated from *Pathway Gene Center* are merged into high scoring clusters via a random walk. To improve candidate modules, a local search is performed in which individual genes are removed, added, or swapped and the module score is returned. Modules that both satisfy the mentioned constraints and result in an increased module score following local search are produced. The user may independently run several iterations of *Clustering* with varied parameters after a single completed execution of *Pathway Gene Center* to compare multiple candidate modules.

*Parameter selection*

Parameters may be modified during the *Clustering* phase. The minimum (-l) and maximum (-u) size of the constructed module can be specified. We recommend varying the minimum average co-expression of the module (-avgCoExpr, recommended range: 0.425-0.52) and the minimum PPI density of the modules (-avgDensity, recommended range: 0.085-0.14). For seeds with generally low pairwise co-expression values, -avgCoExpr can be further reduced. The parameter (-i) is simply an integer used for the initialization of a random number generator. In practice, the number of deleterious loss of function mutations allowed in genes in the module (-m = 6), the minimum ratio of seed scores allowed (-a = 0.5), and minimum pairwise co-expression value allowed (-minCoExpr = 0.01) are not varied.

**Supplementary data**

The Cell-type Specific Expression Analysis (CSEA), Specific Expression Analysis (SEA), and Tissue Specific Expression Analysis (TSEA), and Enrichr tools (Xu *et al.*, 2014; Kuleshov *et al.*, 2016) were applied to each of the 6 modules constructed with pairs of seed genes (CHD8-CREBBP, CHD8-CTNNB1, GABRA3-GABRB1, GRIN2A-GRIN2B, SCN1A-SCN2A, and SHANK2-SHANK3). Enriched KEGG, Gene Ontology (GO) Biological Process, and Online Mendelian Inheritance in Man (OMIM) Expanded terms are displayed for each of the 6 modules in **Supplementary Table 1**. If significant selective expression was detected via CSEA, SEA, and or TSEA for the provided module, respective selective expression plots are displayed in **Supplementary Figure 1**.

### Supplementary Figures

#### CHD8-CREBBP (-min)

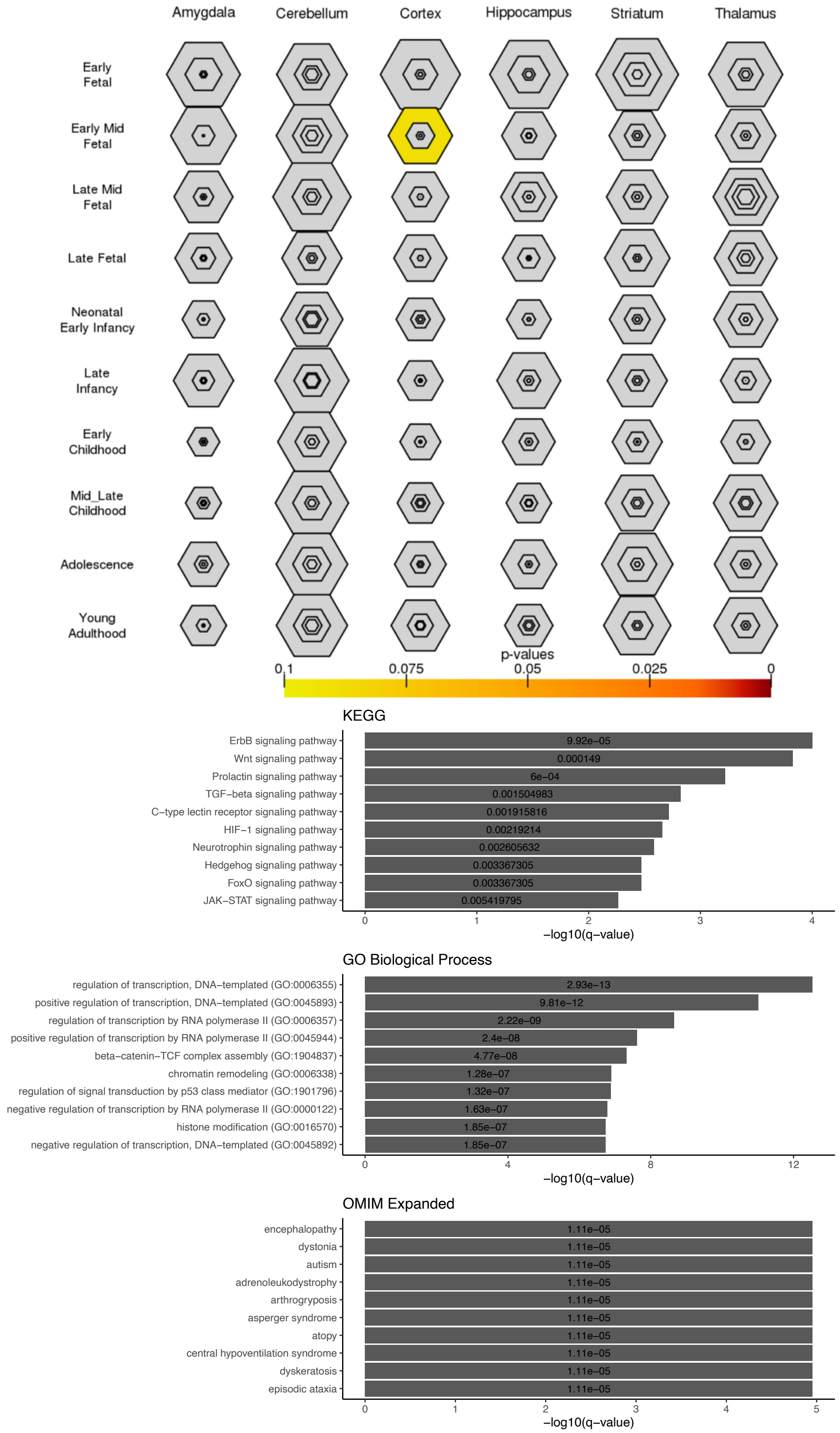

### CHD8-CREBBP (-avg)

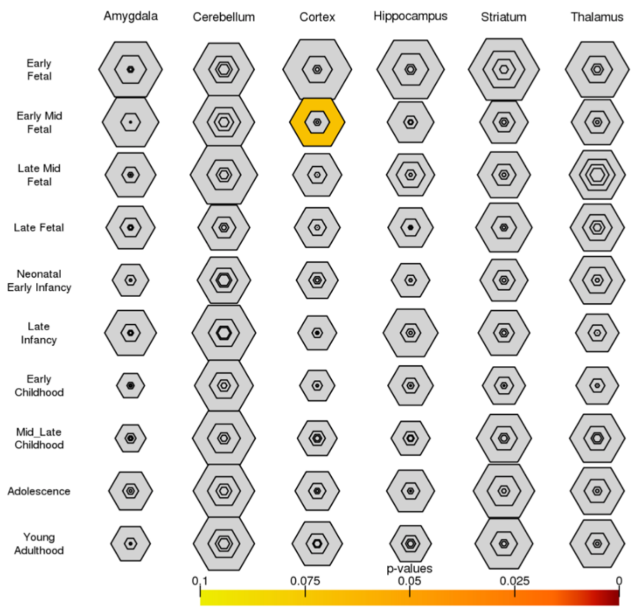


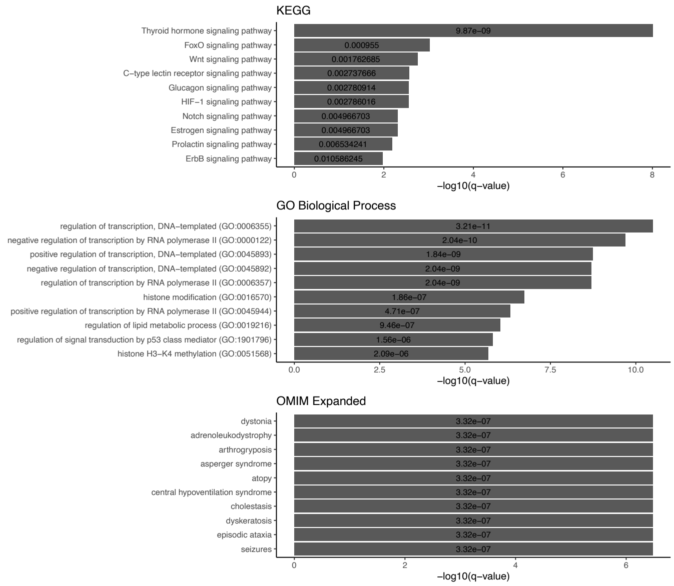


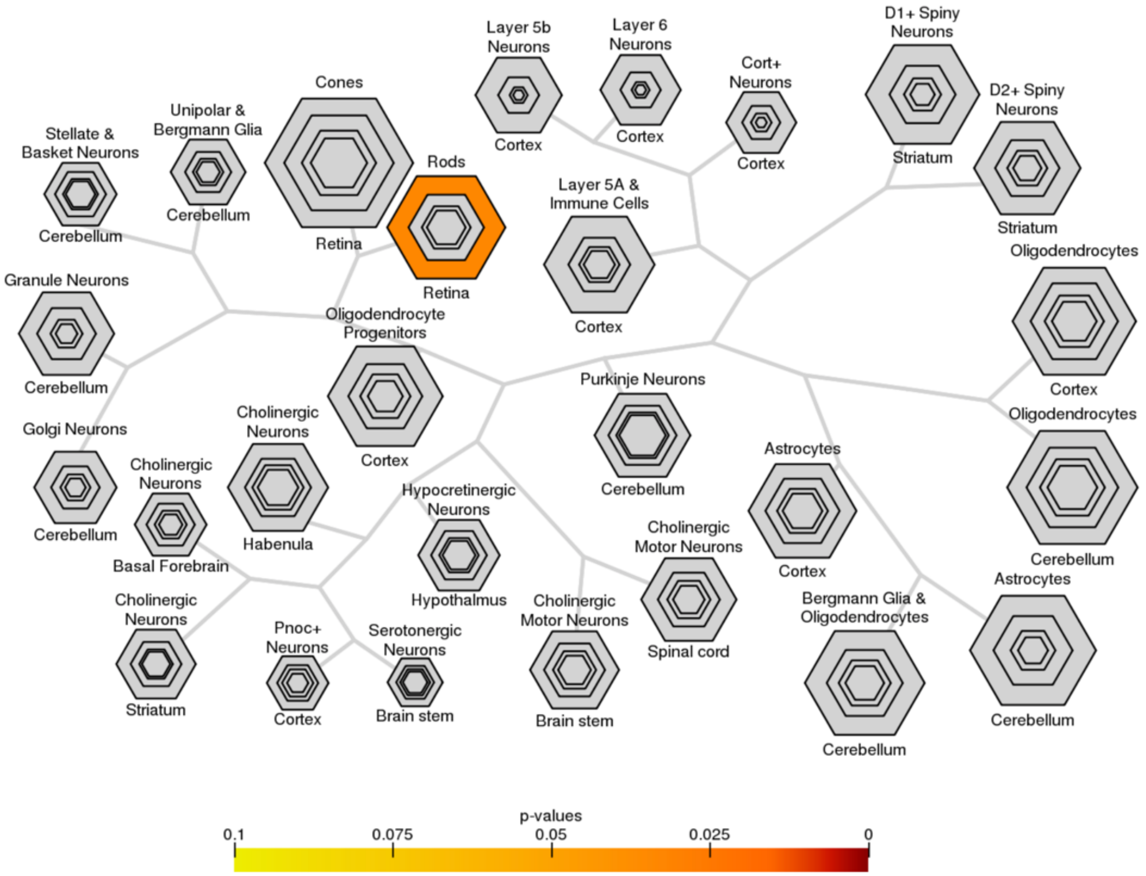

### CHD8-CTNNB1 (-min)

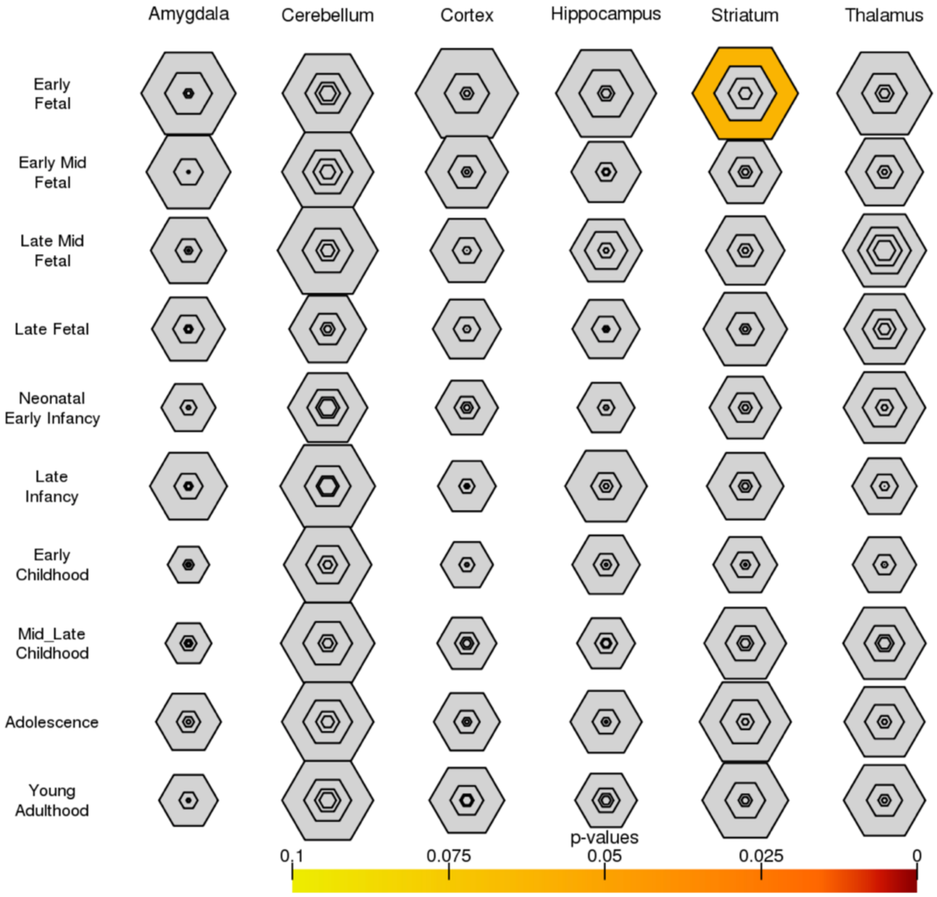


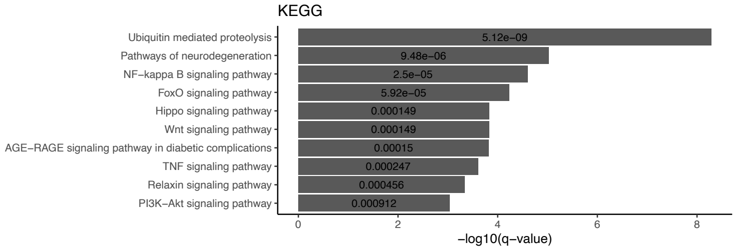


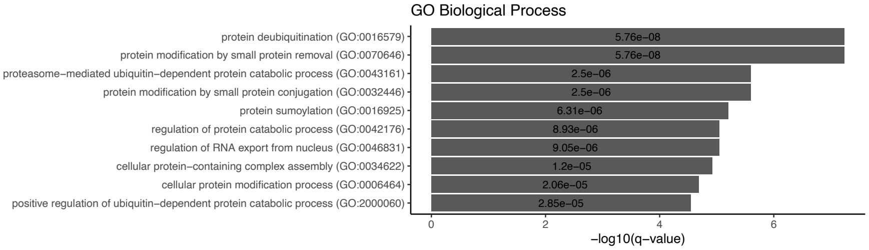


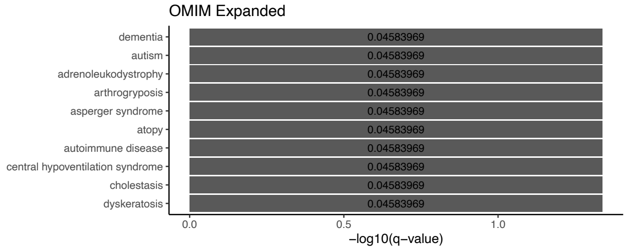

**CHD8-CTNNB1 (-avg)**

No significant enrichment via CSEA, SEA, or TSEA tools.

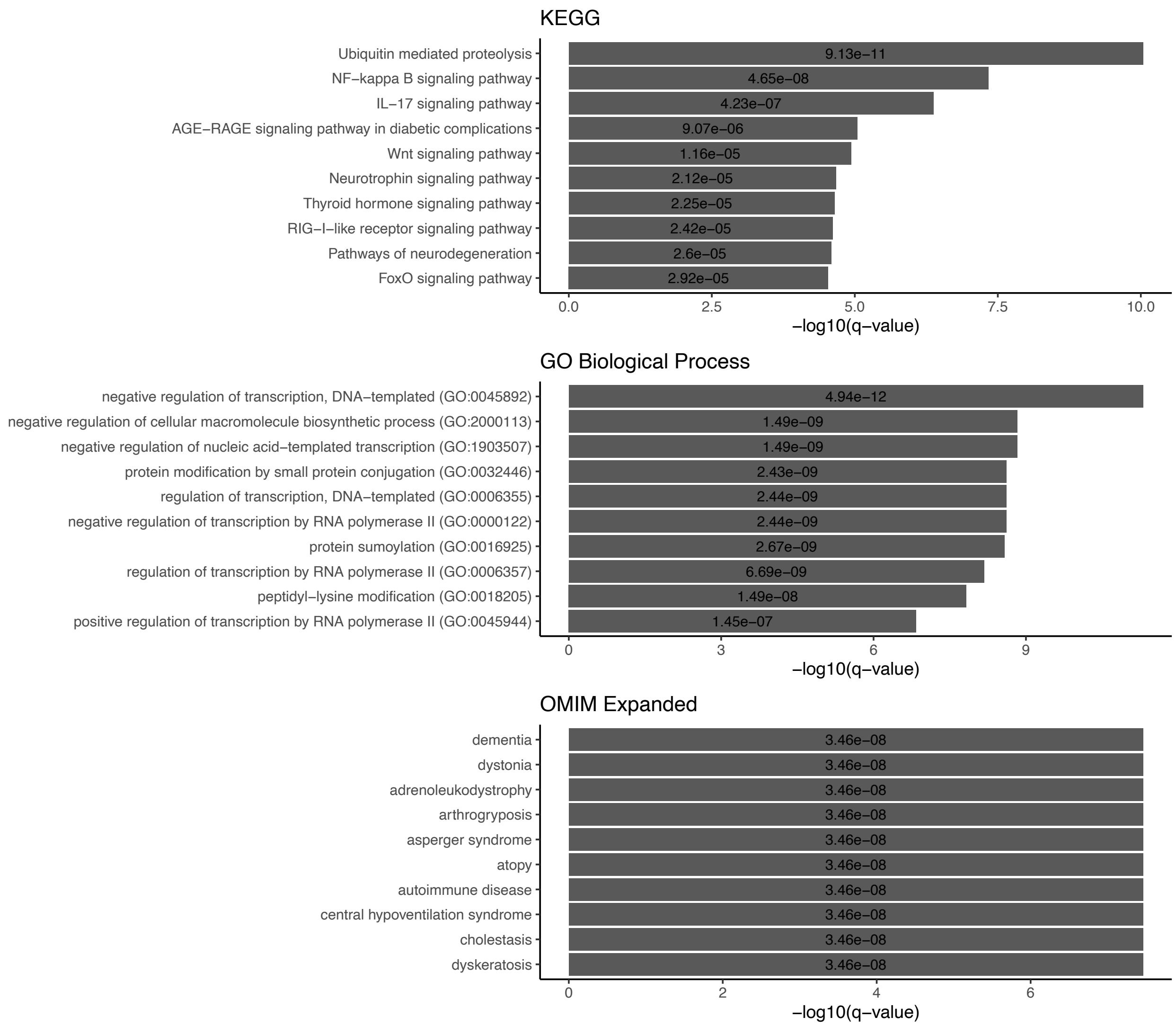

### GABRA3-GABRB1 (-min)

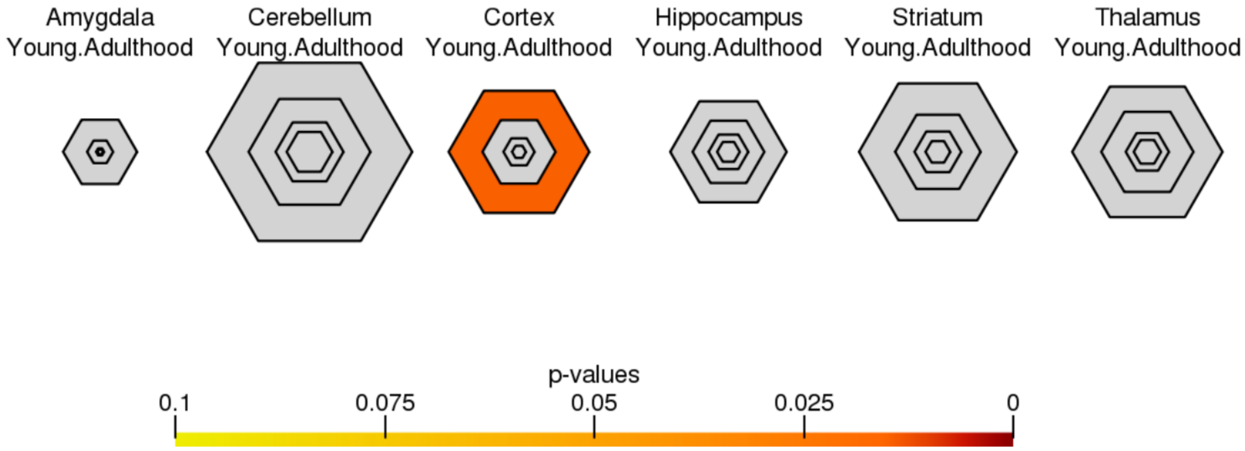


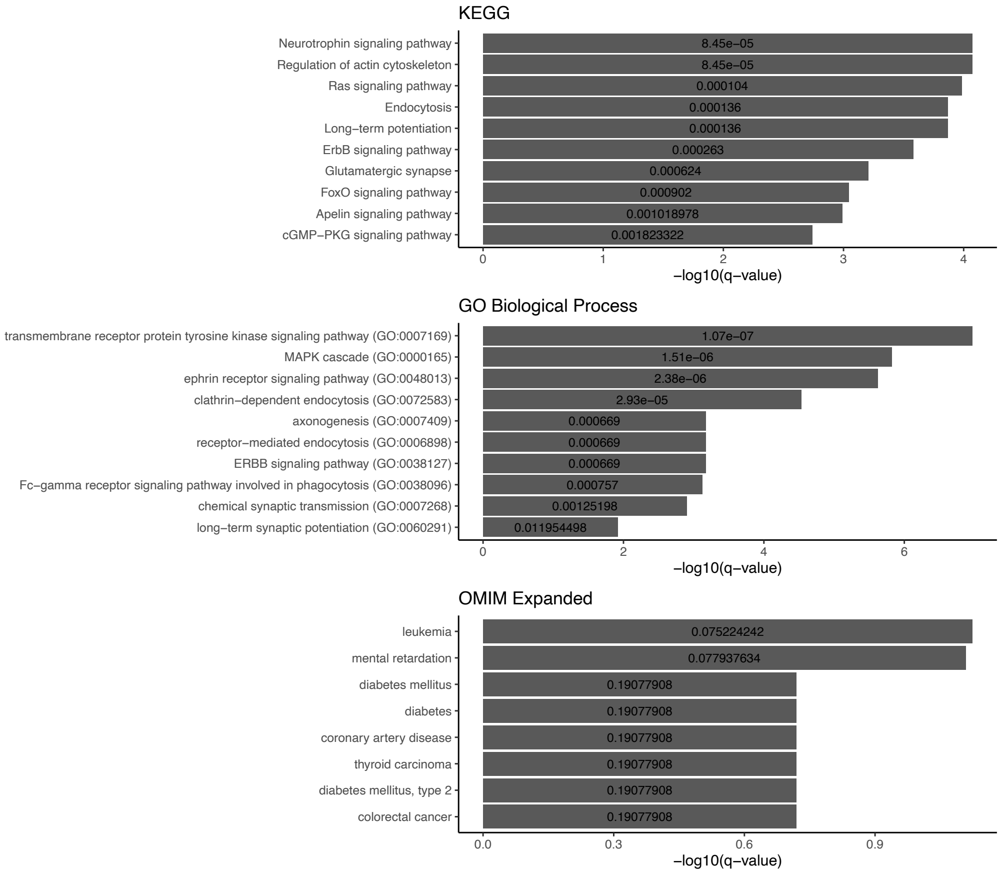

### GABRA3-GABRB1 (-avg)

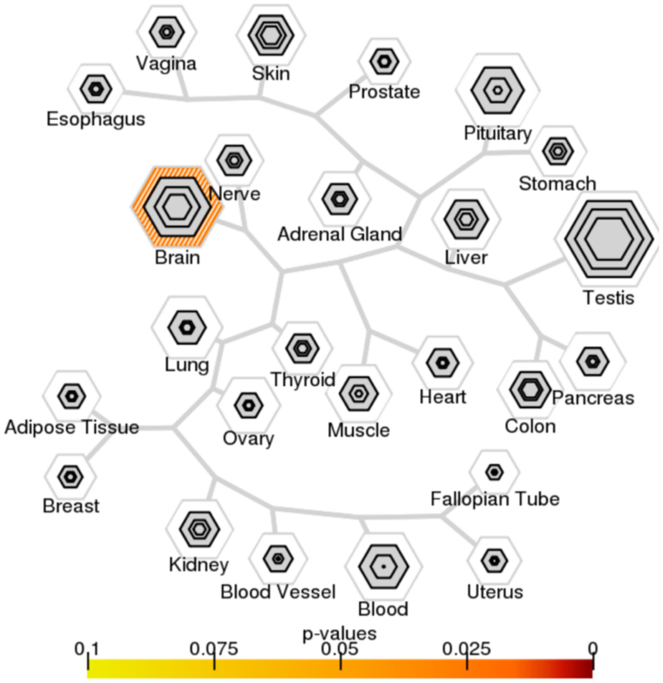


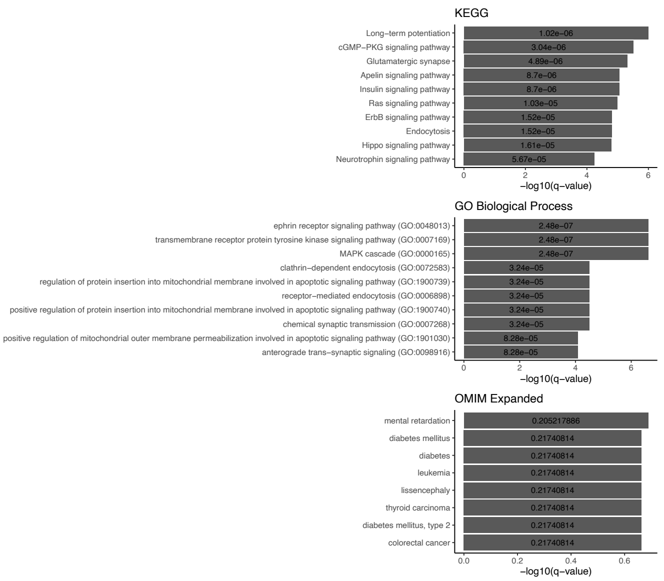


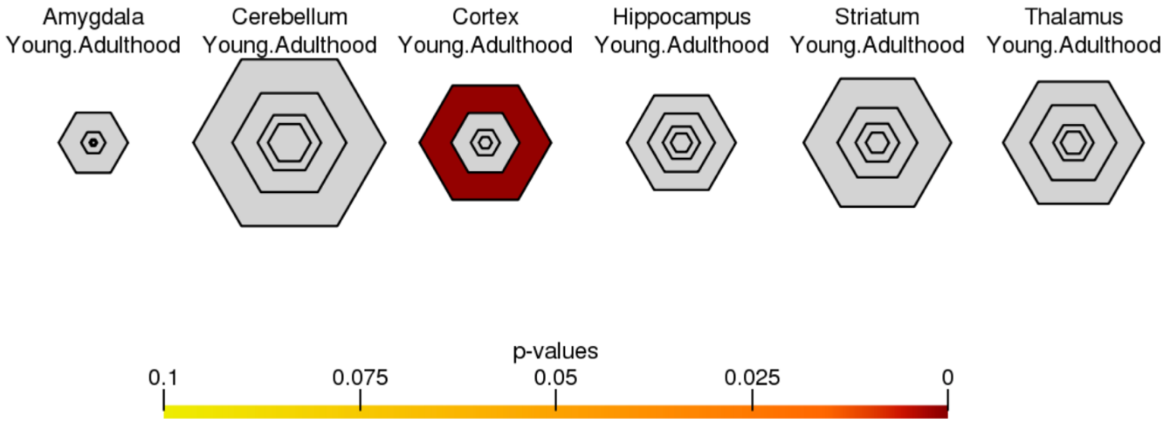

### GRIN2A-GRIN2B (-min)

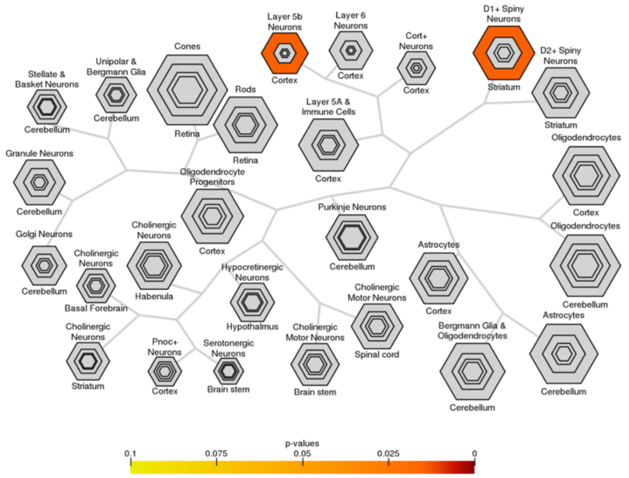


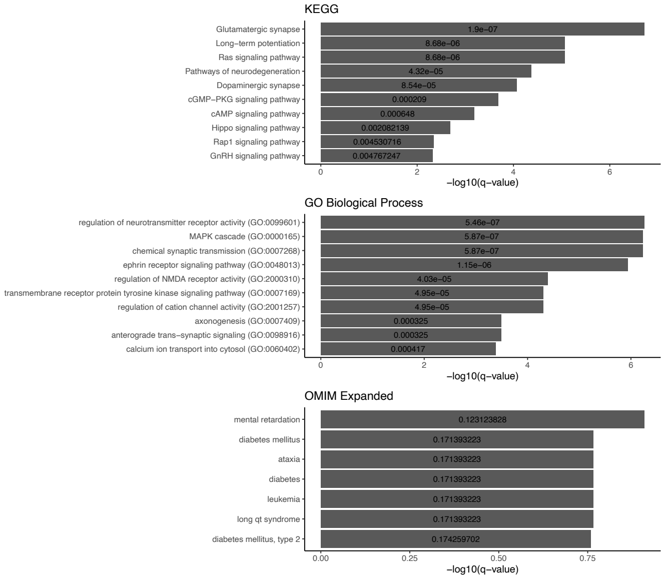


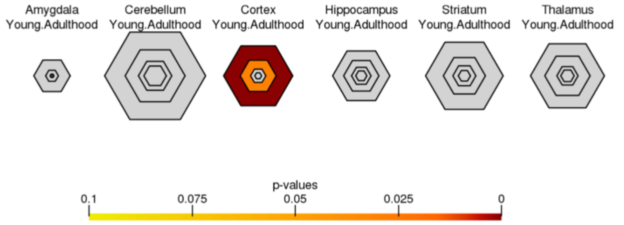


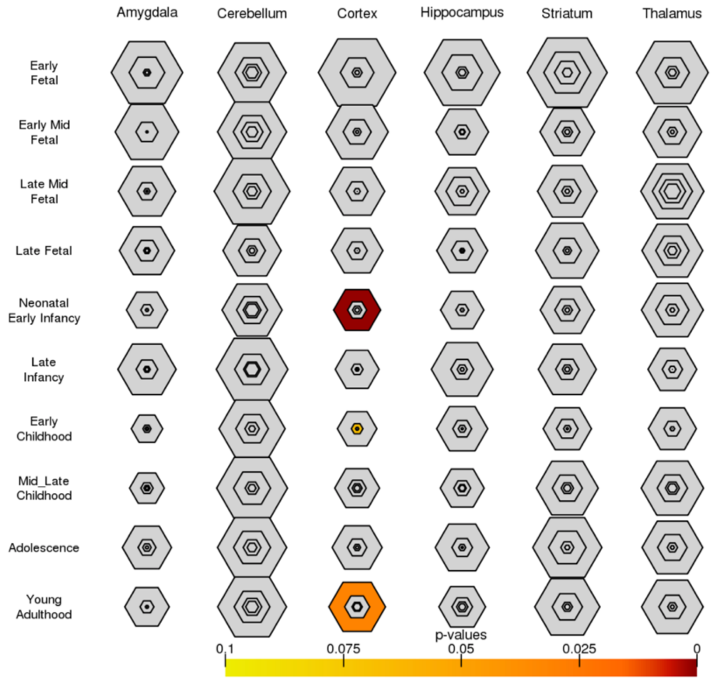


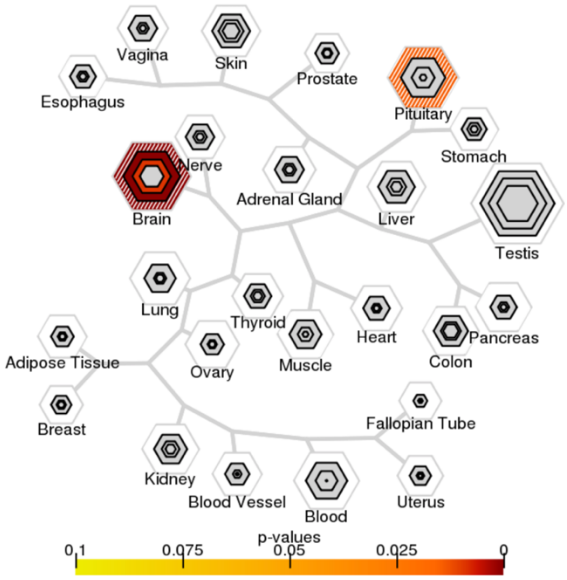

### GRIN2A-GRIN2B (-avg)

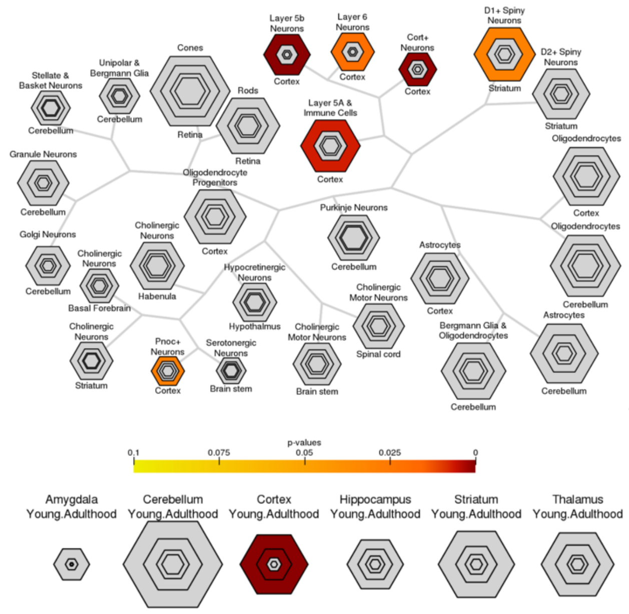


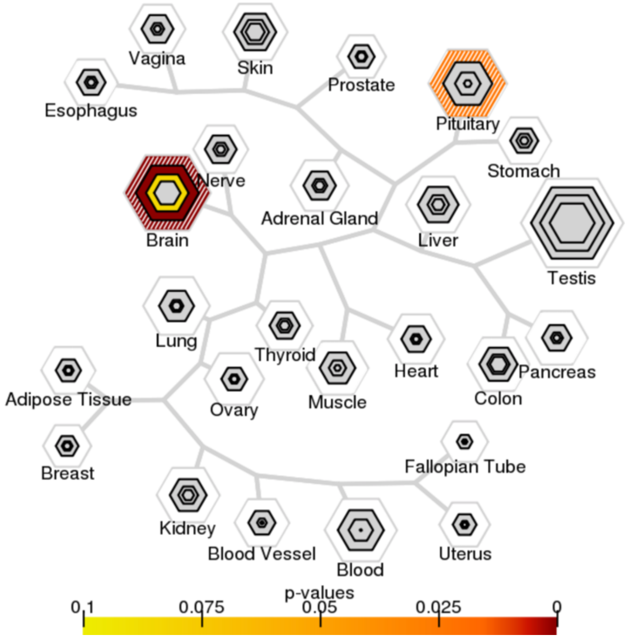


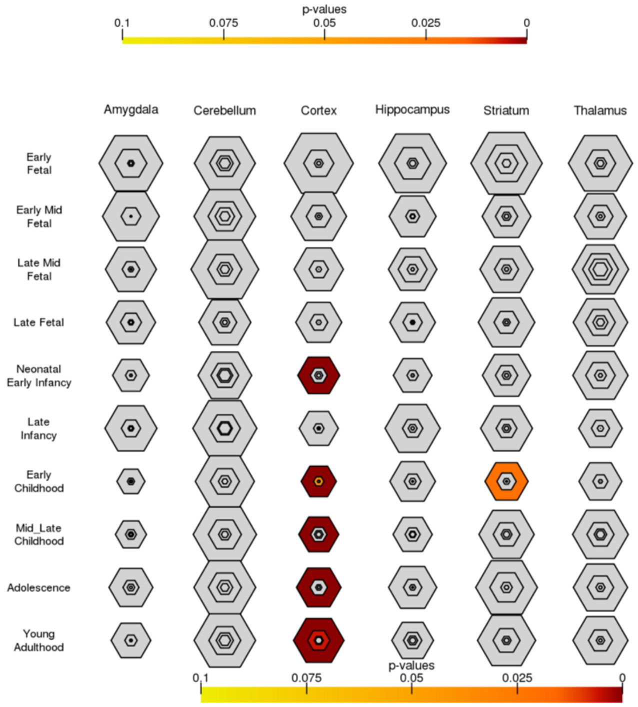


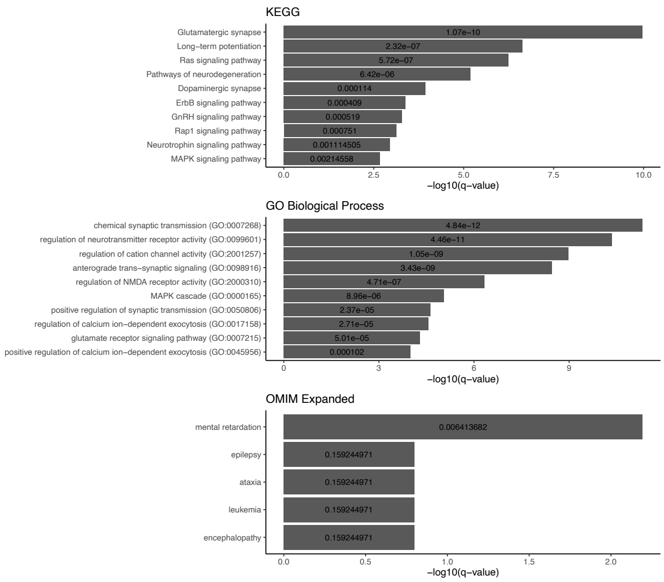

### SCN1A-SCN2A (-min)

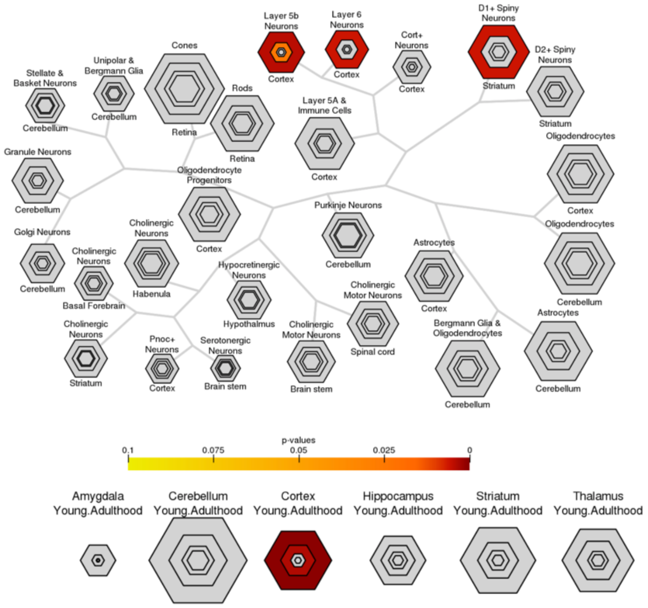


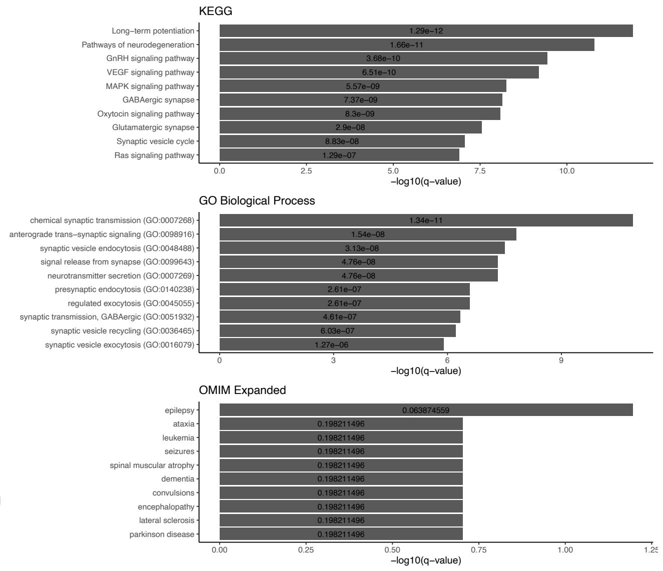


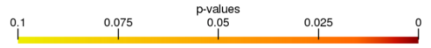


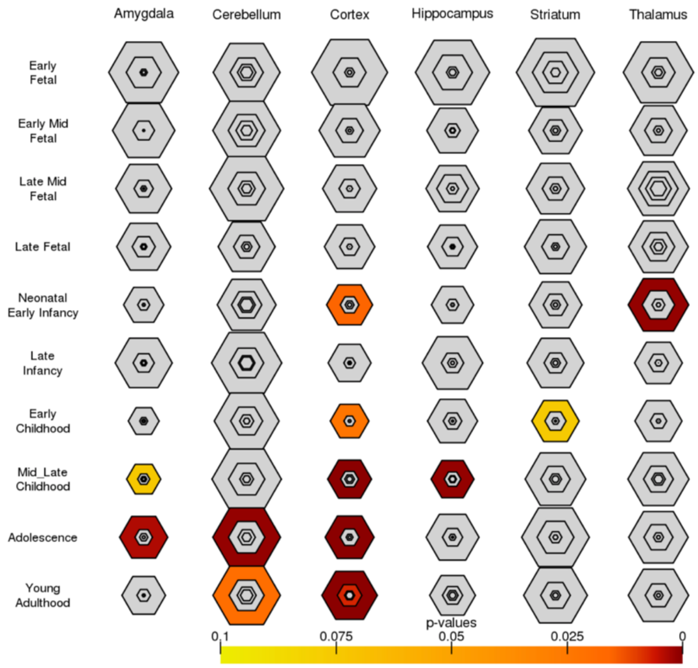


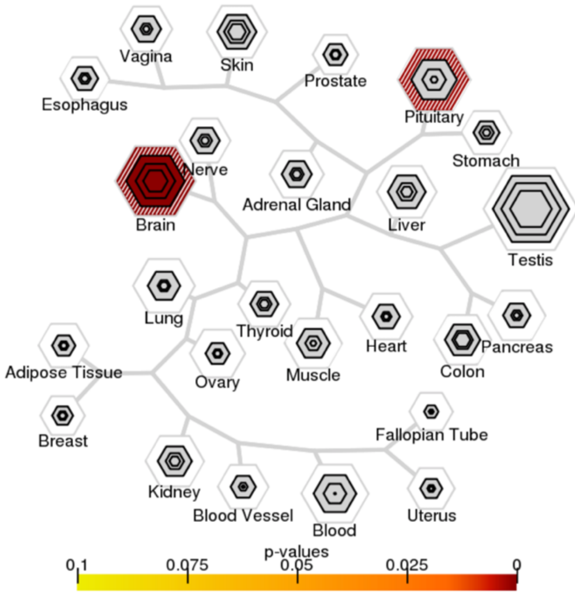

### SCN1A-SCN2A (-avg)

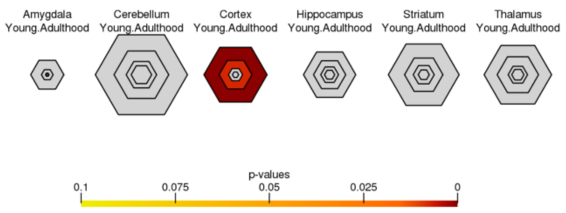


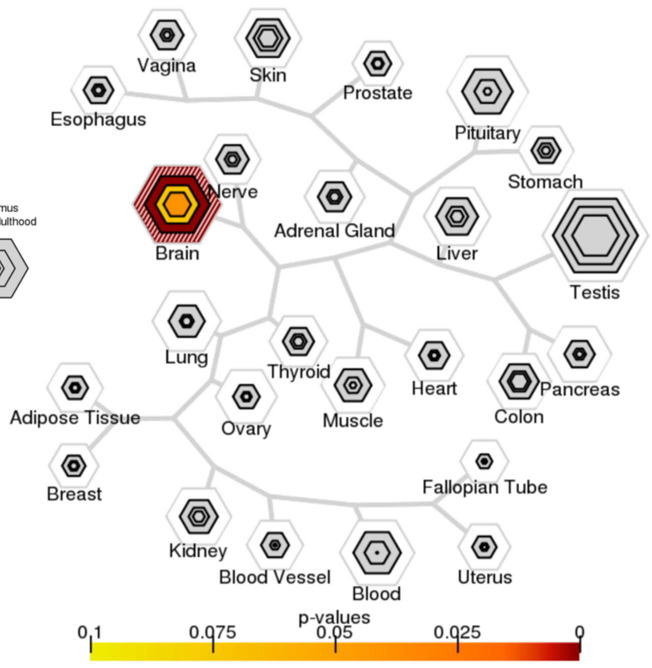


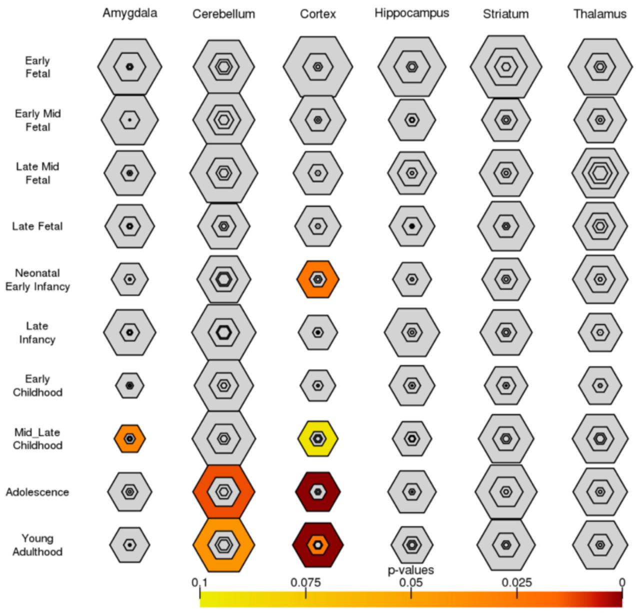


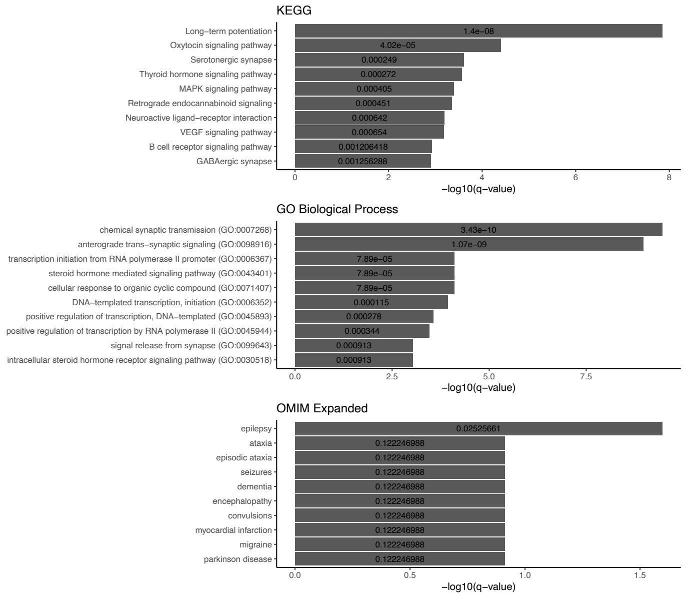

### SHANK2-SHANK3 (-min)

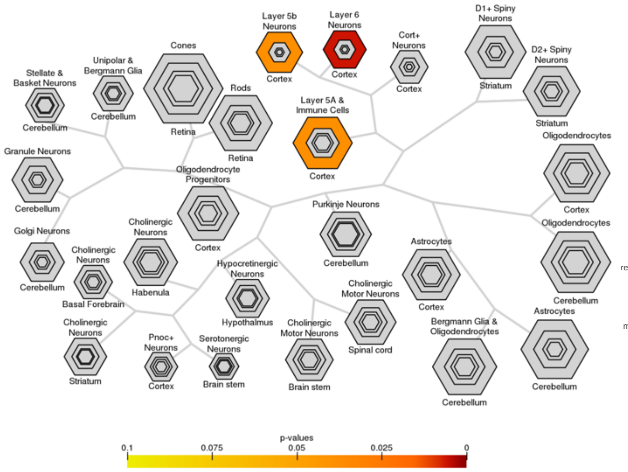


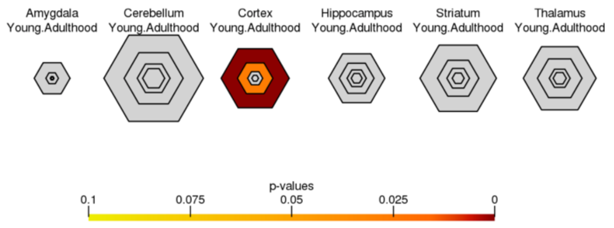


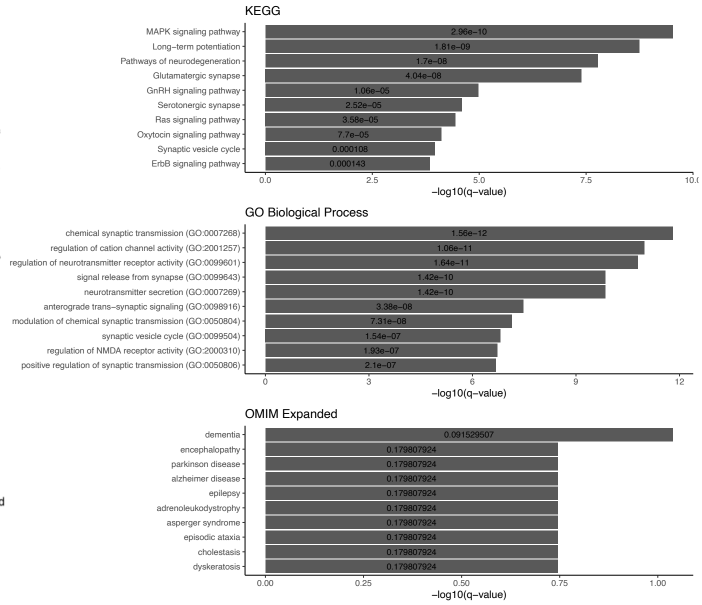


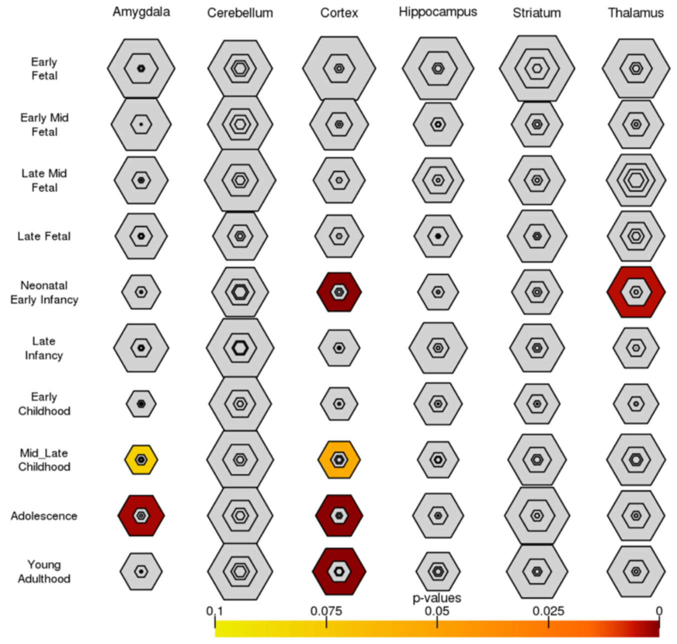


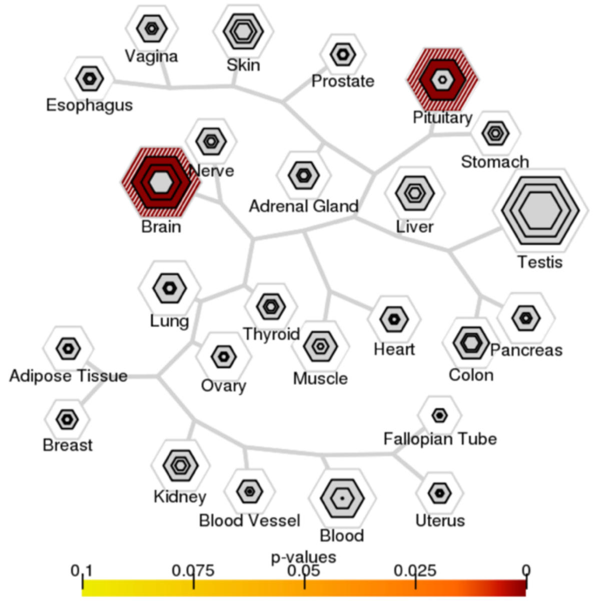

### SHANK2-SHANK3 (-avg)

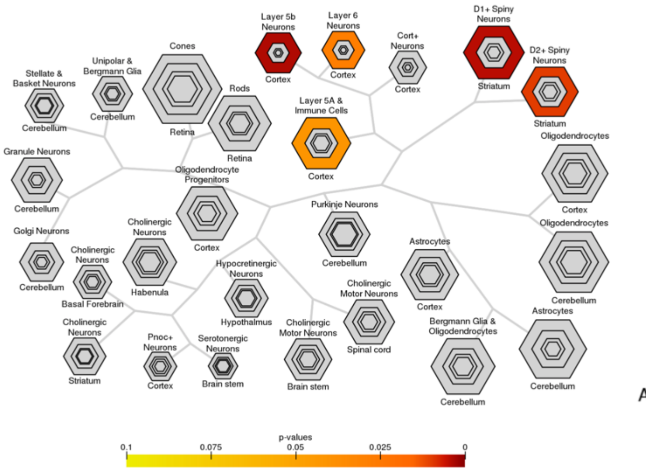


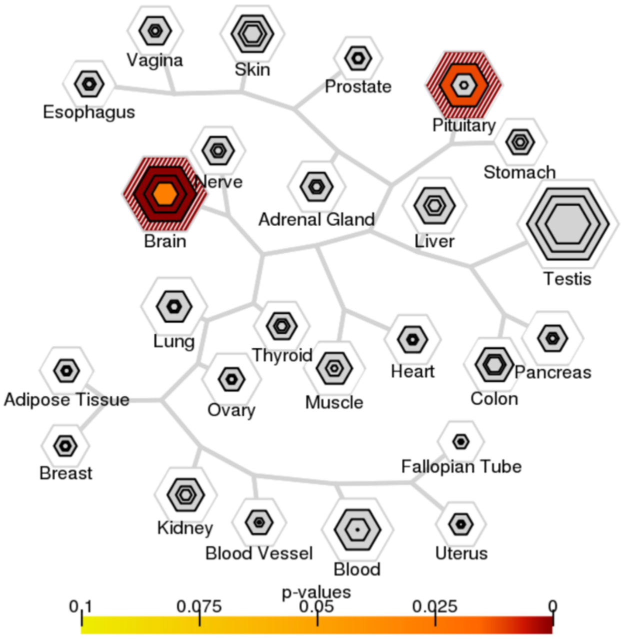


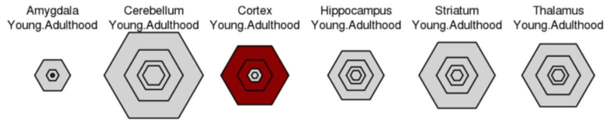


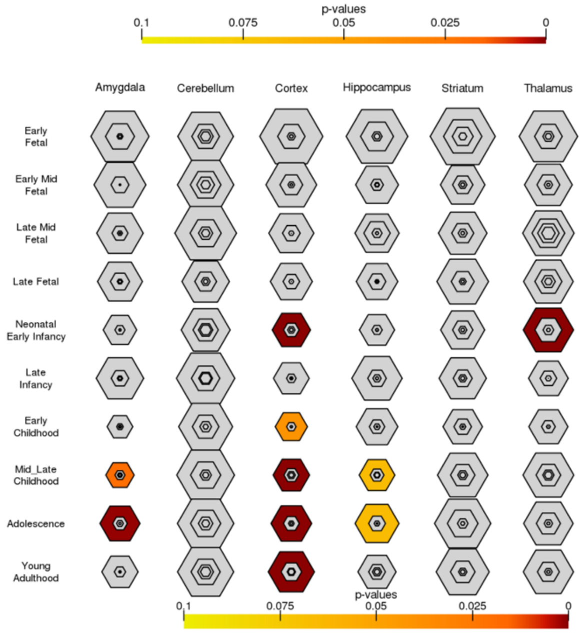


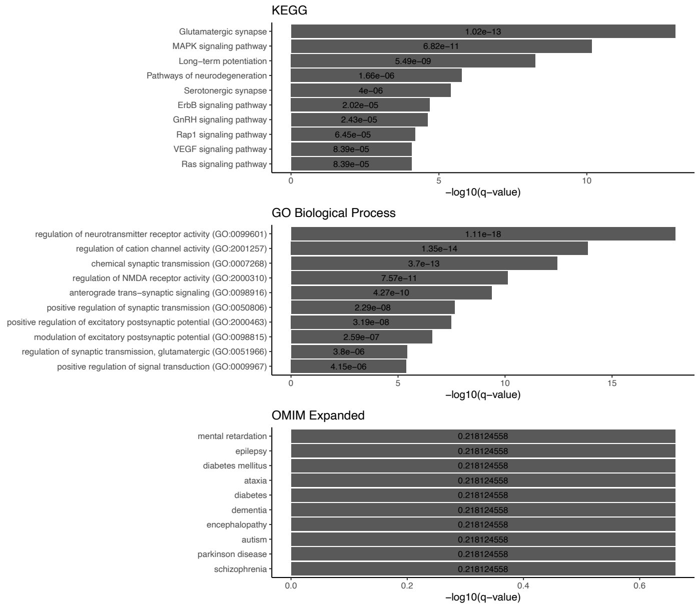

**Supplementary Figure 1.** Significant cell-type and tissue specific expression analysis. If significant selective expression is detected via the CSEA, SEA, and TSEA tools for a module generated using either average (-avg) or minimum (-min) co-expression values during score assignment in *Pathway Gene Center*, corresponding figures are shown. CSEA and SEA describe the significant enrichment of provided genes across cell-types in the human brain at various developmental time periods. TSEA identifies over-representation of provided (disease) genes with enriched expression in certain human tissues. Bar charts display selected functional enrichment terms (**Supplementary Table 1**). Corresponding adjusted p-value (q-value) labels are shown within individual bars.

#### References


Alon,N. *et al.* (1995) Color-coding. *J. ACM*, **42**, 844–856.

Chow,J. *et al.* (2019) Dissecting the genetic basis of comorbid epilepsy phenotypes in neurodevelopmental disorders. *Genome Med.*, **11**, 65.

Hormozdiari,F. *et al.* (2015) The discovery of integrated gene networks for autism and related disorders. *Genome Res.*, **25**, 142–154.

Keshava Prasad,T.S. *et al.* (2009) Human Protein Reference Database—2009 update. *Nucleic Acids Res.*, **37**, D767–D772.

Kuleshov,M.V. *et al.* (2016) Enrichr: a comprehensive gene set enrichment analysis web server 2016 update. *Nucleic Acids Res.*, **44**, W90–W97.

Miller,J.A. *et al.* (2014) Transcriptional landscape of the prenatal human brain. *Nature*, **508**, 199–206.

Szklarczyk,D. *et al.* (2011) The STRING database in 2011: functional interaction networks of proteins, globally integrated and scored. *Nucleic Acids Res.*, **39**, D561–D568.

Xu,X. *et al.* (2014) Cell Type-Specific Expression Analysis to Identify Putative Cellular Mechanisms for Neurogenetic Disorders. *J. Neurosci.*, **34**, 1420–1431.
